# The two zinc fingers in the nucleocapsid domain of the HIV-1 Gag precursor are equivalent for the interaction with the genomic RNA in the cytoplasm, but not for the recruitment of the complexes at the plasma membrane

**DOI:** 10.1101/2020.01.24.918508

**Authors:** E. Boutant, J. Bonzi, H. Anton, M. B. Nasim, R. Cathagne, E. Réal, D. Dujardin, P. Carl, P. Didier, J-C. Paillart, R. Marquet, Y. Mély, H. de Rocquigny, S. Bernacchi

## Abstract

The HIV-1 Gag precursor specifically selects the unspliced viral genomic RNA (gRNA) from the bulk of cellular and spliced viral RNAs *via* its nucleocapsid (NC) domain and drives gRNA encapsidation at the plasma membrane (PM). To further identify the determinants governing the intracellular trafficking of Gag-gRNA complexes and their accumulation at the PM, we compared, in living and fixed cells, the interactions between gRNA and wild-type (WT) Gag or Gag mutants carrying deletions in NC zinc fingers (ZFs), or a non-myristoylated version of Gag. Our data showed that the deletion of both ZFs simultaneously or the complete NC domain completely abolished intracytoplasmic Gag-gRNA interactions. Deletion of either ZF delayed the delivery of gRNA to the PM but did not prevent Gag-gRNA interactions in the cytoplasm, indicating that the two ZFs display redundant roles in this respect. However, ZF2 played a more prominent role than ZF1 in the accumulation of the ribonucleoprotein complexes at the PM. Finally, the myristate group which is mandatory for anchoring the complexes at the MP, was found to be dispensable for the association of Gag with the gRNA in the cytosol.

**STATEMENT of SIGNIFICANCE:** Formation of HIV-1 retroviral particles relies on specific interactions between the retroviral Gag precursor and the unspliced genomic RNA (gRNA). During the late phase of replication, Gag orchestrates the assembly of newly formed viruses at the plasma membrane (PM). It has been shown that the intracellular HIV-1 gRNA recognition is governed by the two-zinc finger (ZF) motifs of the nucleocapsid (NC) domain in Gag. Here we provided a clear picture of the role of ZFs in the cellular trafficking of Gag-gRNA complexes to the PM by showing that either ZF was sufficient to efficiently promote these interactions in the cytoplasm, while interestingly, ZF2 played a more prominent role in the relocation of these ribonucleoprotein complexes at the PM assembly sites.

## INTRODUCTION

During the late phase of human immunodeficiency virus type-1 (HIV-1) replication, the retroviral 55-kDa precursor (Pr55^Gag^ or Gag) orchestrates the assembly of newly formed viruses at the plasma membrane (PM) (1–3). Gag specifically selects the HIV-1 unspliced genomic RNA (gRNA) from the bulk of cellular and spliced viral RNAs, for encapsidation *via* its nucleocapsid domain (NC). This process involves specific interactions between Gag and the 5’ end of the gRNA which contains the packaging signal (Psi) encompassing stem-loop 1 (SL1) to SL4 (Figure 1A) (for reviews see (4–7)). SL1 corresponds to the Dimerization Initiation Site (DIS), as it contains a short palindromic sequence in its apical loop that drives dimerization of the HIV-1 gRNA (8–12), and our group previously showed that the SL1 internal purine rich loop corresponds to a major Gag recognition signal (13–15). SL2 contains the major splice donor site, SL3 has been historically considered as the main packaging signal (16, 17), and SL4 contains the translation initiation codon of Gag.

**FIGURE 1:**
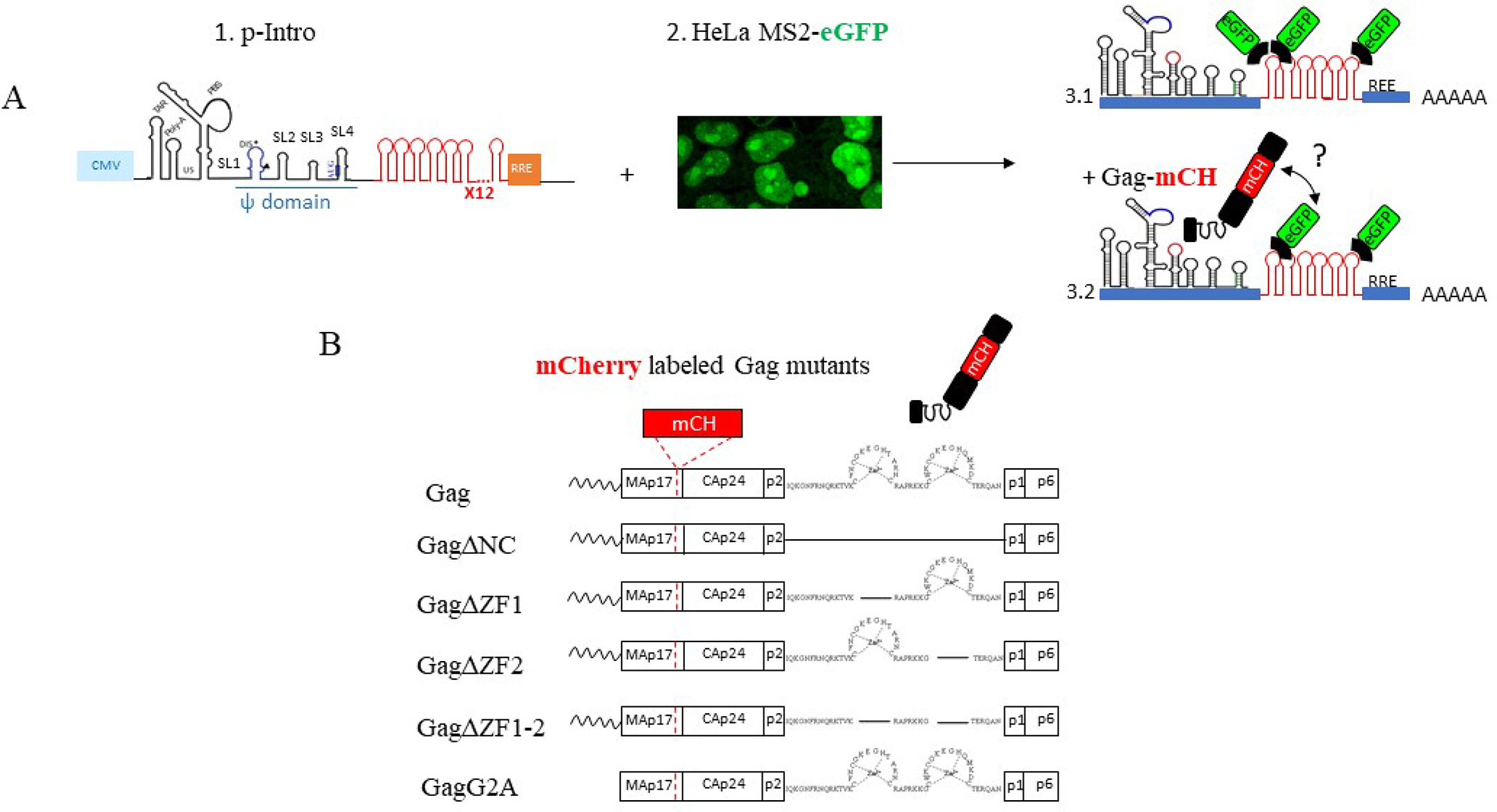
Fluorescent tools used for microscopy imaging. (A) 1. Schematic presentation of the HIV-1 reporter construct (p-Intro). The RNA obtained from the CMV-dependent (light blue square) transcription of this plasmid contains twelve copies of MS2 stem loops (SL) inserted between the ψ domain and RRE element. 2. HeLa cells constitutively expressing MS2-eGFP. The fluorescence was found to be localized in the nucleus and concentrated in the nucleoli in non-transfected cells (23). 3. Schemes of the unspliced nascent HIV-1 mRNAs harbouring the SL elements that can be recognized by dimers of the fluorescently labelled MS2 capsid proteins (MS2-eGFP) and interact with mCH-labelled Gag proteins. eGFP and m-Cherry constitute a donor-acceptor couple for FRET. (B) Schematic representation of the Gag proteins used in this study. Gag domains are represented: from N-terminus the myristoyl group, MA (Matrix), CA (Capsid), NC (Nucleocapsid) including two zinc fingers (ZFs), the spacer peptides p1 and p2 and the C-terminal p6 domain. We deleted either the entire NC domain (GagΔNC), or both ZFs (GagΔZF1-2) or only one ZF (GagΔZF1 and GagΔZF2). We also included in our study a non-myristoylated version of Gag (GagG2A). The deletions were represented by a straight line linking the bordering amino acid residues. All the Gag proteins were fused to mCherry (mCH), which was inserted between the MA and CA domains (63).

Using imaging techniques, several groups showed that gRNA dimerization precedes the budding of viral particles (18–23). Indeed, HIV-1 gRNA was found to migrate to the PM as a pre-formed dimer (18, 23, 24) in association with low-order Gag multimers (25–27), thus forming a viral ribonucleoprotein (vRNP) (for reviews (6, 28, 29)). Although the cellular trafficking of the vRNP remains to be precisely described, it was proposed that the viral core was alternatively targeted to late endosomes (30–33), and the dynein motor function could regulate the vRNP egress on endosomal membranes thus impacting viral production (34). Moreover, it was suggested that stabilization of the RNA dimers occurs at the PM assembly sites (24) thanks to the chaperone activity of Gag (18, 19, 25, 26).

The Gag precursor is composed of four main domains and two short spacers (for review see (35)) (Figure 1B), starting from the N-terminus with the matrix (MA) domain, which mediates the association of Gag with the PM (36) *via* its N-terminal myristoylated glycine (G2) and a highly basic region (HBR) which associates with PI(4, 5)P_2_ (37, 38). The capsid (CA) domain drives Gag multimerization, leading to the formation of the structural viral core. Recent studies indicated that CA interacts with MA stabilizing the compact conformation of the precursor (39). The NC domain flanked by two spacer peptides p2 and p1, contains two CCHC ZF motifs and constitutes the major determinant for gRNA recognition (40–42). Importantly, the NC domain was also found to facilitate Gag multimerization and viral assembly (43–46), and its fully matured form, NCp7, fulfills multiple functions in the early steps of the viral cycle by acting as a nucleic acid chaperone. As such, NCp7 is thought to mediate structural gRNA rearrangements (for reviews see (47–49)). Finally, at the C-terminus, the p6 domain promotes the budding of nascent virions at the PM, by interacting with host factors associated with the ESCRT (Endosomal Sorting Complex Required for Transport) machinery (for a review see (50)). Our group recently showed that p6 is also a key determinant for specific Gag-gRNA interaction (51).

Both MA and NC possess nucleic acid (NA) binding properties. MA interacts with NAs *in vitro* and in cells (27, 52) *via* its HBR domain. In particular, its interaction with host tRNAs in the cytosol might regulate Gag interaction with the PM (27). On the other hand, interactions of NC with gRNA are mainly driven by the two highly conserved ZFs (40, 53–58). However, these ZFs do not seem to be functionally equivalent, the N-terminal ZF (ZF1) playing a more prominent role in gRNA selection and packaging (57). Indeed, mutations in ZF1 and the NC N-terminal domain led to formation of particles with abnormal core morphology and affected proviral DNA synthesis (59, 60). Besides, specific *in vitro* binding of NCp7 to the Psi was also found to be dependent on the ZF1 and flanking basic amino acid residues (61). However, the exact contribution of each ZF remains controversial since a recent *in vitro* study showed that the distal C-terminal ZF (ZF2) would drive the first steps of association with NAs, because of its larger accessibility compared to ZF1 which would contribute to stabilize the resulting complex (62).

Here, to decipher the role of the two ZFs in the cellular trafficking of Gag-gRNA complexes to the PM, we combined several quantitative approaches including confocal microscopy, time-lapse microscopy, FRET-FLIM (Fluorescence Resonance Energy Transfer-Fluorescence Lifetime Imaging Microscopy), and RICS (Raster Image Correlation Spectroscopy). To this aim, the MS2 bacteriophage coat protein was fused to eGFP to fluorescently label the gRNA (Figure 1A), whereas Gag proteins were fused to the fluorescent protein probe mCherry (mCH) (Figure 1B). This allowed us to compare in the cytoplasm and at the PM, the interactions of gRNA with WT Gag, and Gag mutants carrying deletions in NC zinc fingers (ZFs) and a non-myristoylated version of Gag, (GagG2A) (Figure 1B). As expected, the GagG2A mutation prevented co-localization of Gag with the gRNA at the PM but did not impair its gRNA binding. Importantly, we found that the simultaneous deletion of the two ZFs completely abolishes the Gag-gRNA interactions in the cytosol and at the PM. Either ZF was found to be sufficient to efficiently promote Gag-gRNA interactions in the cytoplasm, hence displaying redundant roles in this respect. Interestingly, ZF2 played a more prominent role than ZF1 in the relocation of these ribonucleoprotein complexes at the PM. Taken together, we show here that the intracellular HIV-1 gRNA recognition and Gag-gRNA trafficking to the PM are governed by ZF motifs within the NC domain.

## MATERIALS and METHODS

### Plasmids DNA

The constructs for Gag, and Gag-mCherry (Gag-mCH) were previously described (44, 63). The plasmid encoding human-codon-optimized Gag was kindly provided by David E. Ott (National Cancer Institute at Frederick, Maryland). The deletion mutants (GagΔNC, GagΔZF1, GagΔZF2 and GagΔZF1-2) and the substitution mutant (GagG2A) were constructed by PCR-based mutagenesis on Gag, and Gag-mCH following the supplier’s protocol (Stratagene). In addition, the p-Intro plasmid was obtained from E. Bertrand (IGM, Montpellier, (30)) and modified by Epigex (Strasbourg – [pcDNA3.1 plasmid; CMV promotor]). Then, a TAG codon was introduced in the plasmid to stop the expression of eCFP-SKL (peroxisome localization signal) using Phusion site-directed mutagenesis kit (Thermo scientific F-541) and a set of primers [FW: 5’-GATATGGTGAGCTAGGGCGAGGAGCTG-3’ and Rev: 5’-GATACCGTCGAGATCCGTTCACTAATCG-3’]. The plasmid pPOM21-mCH was obtained from Euroscarf (64), while pRSV-Rev was obtained from Addgene (Plasmid: ♯12253). The integrity of all plasmids was assessed by DNA sequencing (GATC Eurofins Genomics).

### Cell culture and plasmid transfection

HeLa cells stably expressing homogenous levels of MS2-GFP with a nuclear localization signal (so called MS2-GFP) were obtained from Nolwenn Jouvenet (Institute Pasteur, Paris) (23) and grown in Dulbecco’s modified Eagle medium (DMEM, Gibco LifeTechnology 11880-028) supplemented with 10% fetal bovine serum (FBS, Lonza), 1% antibiotic solution (penicillin-streptomycin, Gibco-Invitrogen) and glutamine at 37°C in humidified atmosphere containing 5% CO_2_.

To study the interaction between gRNA and Gag *in cellula*, MS2-eGFP HeLa cells were seeded onto a cover glass in 12-well plates (see confocal and super resolution experiments) or onto an IBIDI® chambered cover glass (see FRET-FLIM experiments) at the density of 7.5×10^4^ cells/mL/well or 1.5×10^5^ cells/mL/well, respectively, 24 h before transfection. MS2-eGFP HeLa cells were then transfected using JetPrime™ (Life Technologies, Saint Aubin, France) with a mixture of plasmids encoding for pIntro, Rev, unlabeled Gag and Gag -mCH proteins at the following ratios depending of the material used [12 well plate: 1; 0.25; 0.2 µg – in IBIDI® chamber: 1.6;0.4;0.1 µg]

### Immunolabeling

The MS2-eGFP HeLa cells were fixed 24 h post-transfection with 1.5-4% of Paraformaldehyde PFA/PBS for 15 min and then rinsed three times for 5 min with PBS. Cells were then permeabilized with 0.2% Triton X-100, blocked in 3% (W/V) BSA for 1 h, and subsequently incubated for 1 h at room temperature with rabbit polyclonal antibody directed against RNA polymerase II phosphoS2 (Abcam - ab5095), followed by an incubation with fluorescent Alexa-568 anti-rabbit secondary antibody (ThermoFischer Scientific A11011). For nuclear staining, the medium was replaced by Hoechst 33258 (Molecular Probes, 5 µg/mL) in PBS and cells were incubated for 10 min. Coverslips were then washed and mounted on microscope slides with Fluoromount-G (Thermo Fischer Scientific 00-4958-02). Images were acquired with a Leica TCS SPE II confocal microscope equipped with a 63×1.4 NA oil immersion objective (HXC PL APO 63x/1.40 OIL CS) and 405, 488 and 561 nm laser diodes.

### Confocal microscopy

Fluorescence confocal images of tagged Gag proteins in fixed cells in presence or absence of MS2-eGFP were taken 24 h post-transfection using a Leica SPE microscope equipped with a 63×1.4 NA oil immersion objective (HXC PL APO 63x/1.40 OIL CS). The eGFP images were obtained by scanning the cells with a 488 nm laser line and using a 500-555 nm emission bandwidth. For the mCH images, a 561 nm laser line was used with a 570-625 nm bandwidth filter. To quantify the phenotypes, we first analyzed Gag proteins localized at the membrane (Red channel), and then checked if MS2-eGFP labelled RNA localized at the membrane or in the cytoplasm (Green channel). We assessed 100 cells per experiment, and three independent experiments were performed.

### Fluorescence Lifetime Imaging Microscopy (FLIM)

The experimental set-up for FLIM measurements was previously described (44). Briefly, time-correlated single-photon counting FLIM measurements were performed on a home-made two-photon excitation scanning microscope based on an Olympus IX70 inverted microscope with an Olympus 60 × 1.2 NA water immersion objective operating in the scanned fluorescence collection mode. Two-photon excitation at 900 nm was provided by an Insight Deep see laser (Spectra Physics). Photons were collected using a short pass filter with a cut-off wavelength of 680 nm (F75-680, AHF, Germany) and a band-pass filter of 520 ± 17 nm (F37-520, AHF, Germany). The fluorescence was directed to a fiber coupled APD (SPCM-AQR-14-FC, Perkin Elmer), which was connected to a time-correlated single photon counting module (SPC830, Becker & Hickl, Germany).

The time-resolved decays were analyzed using a one component model pixel *per* pixel to obtain the fluorescence lifetime distribution all over the cell. Numerical values were converted into an arbitrary color scale producing an image ranging from blue (presence of FRET) to yellow (absence of FRET).

For Förster Resonant Energy Transfer (FRET) experiments, the FRET efficiency (E) was calculated according to the equation:

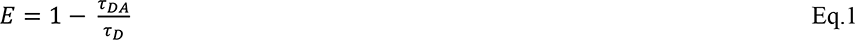

where τ_DA_ and τ_D_ are the lifetime of the donor in the presence and in the absence of the acceptor. To observe gRNA-Gag interactions in live-cells, the seeded cells (on IBIDI® chamber) were transfected as described above and washed once with PBS. A freshly prepared Leibovitz’s L15 Medium (Gibco 21083-027) with FBS was added prior to observation.

### DNA plasmid Microinjection and time-lapse microscopy

For time-lapse experiments, sub confluent MS2-eGFP HeLa cells plated on glass coverslips (in a 12 well plate at 1.5×10^5^ cells/mL the day prior the experiment) were mounted in a Ludin Chamber (Life Imaging Services, Basel, Switzerland) following the protocol described in (44). The cells were then placed on a Leica DMIRE2 microscope equipped with a chamber at 37 °C, with 5% CO_2_ (Life Imaging Services). A mixture of plasmids (72% pIntro, 17% Rev, 5.5% Gag and 5.5% mCH-Gag or Gag mutants in the NC domain) were microinjected into the nucleus at 0.1 μg/μL with a fluorescent microinjection reporter solution (0.5 μg/μL rhodamine dextran; Invitrogen), using a Femtojet/InjectMan NI2 microinjector (Eppendorf). The coordinates of several microinjected cells were memorized using a Märzhäuser (Wetzlar, Germany) automated stage piloted by the Leica FW4000 software. Images were then acquired with a 100 × HCX PL APO (1.4 NA) objective every 5 min during 2-4 h using a Leica DC350FX CCD camera controlled by the FW4000 software. Time-lapse movies were then analyzed using the MetaMorph (Molecular Devices, Sunnyvale, USA) and ImageJ (National Institutes of Health, USA) software in order to determine at which time the GFP signal appears at the PM (fluorescently labeled gRNA), as well as when Gag multimers appear at the PM.

### Raster Image Correlation Spectroscopy (RICS)

MS2-eGFP HeLa cells were transfected with specific plasmids (in Ibidi® Chamber) as described above and living cells were imaged at 16 h post-transfection. Raster Image Correlation Spectroscopy (RICS) measurements were performed on a Leica SPE microscope equipped with a 63 × oil immersion objective (HXC PL APO 63x/1.40 OIL CS Leica). eGFP and mCH were excited with 488 nm and 561 nm laser lines, respectively. The emitted fluorescence was detected by a photo multiplier (PMT) with a detection window of 500-550 nm and 590-700 nm for eGFP and mCH, respectively. For each RICS measurement, a stack of 50 images (256 × 256 pixels with a pixel size of 50 nm) was acquired at 400 Hz (2.5 ms between the lines with a pixel dwell time of 2.8 µs). The RICS analysis was then performed using the SimFCS software developed by the Laboratory of Fluorescence Dynamics (http://www.lfd.uci.edu), or alternatively by a package of plugins running under ImageJ software (https://imagej.nih.gov/ij/). In the latter case, the used tools were an extension and improvement of the Stowers ICS Plugins developed by Jay Unruh (http://research.stowers.org/imagejplugins/ics_plugins.html) allowing us to generate RICS maps over several acquisitions in a fully automated and optimized way.

Before the autocorrelation of the image, the contribution of the slowly moving structures and cellular displacements were removed by subtracting the moving average. Then, the correlations of all frames were calculated, and the final averaged autocorrelation surface was fitted with the RICS correlation function given by:

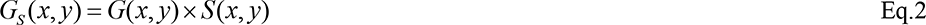

where G(x,y) represents the temporal correlation resulting from the diffusion of the fluorescent molecules and S(x, y) takes into account the effect of beam displacement in the x and y directions. These two terms are defined as:

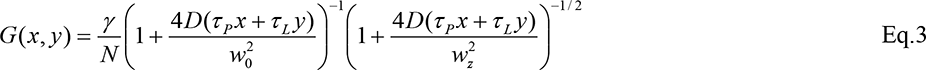

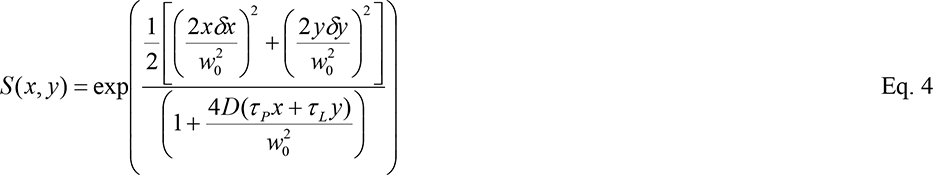

where x and y are the spatial lags in pixels and δx and δy are the pixel size (50 nm). τ_P_ and τ_L_ are the pixel dwell time (2.8 µs) and the interline time (2.5 ms), respectively. w_0_ is the beam waist, w_z_ represents the z-axis beam radius and is set to 3w_0_. γ is a shape factor due to uneven illumination across the focal volume and is 0.3535 for a 3D Gaussian beam. N and D are the floating parameters that represent the number of fluorescent molecules in the focal volume and the diffusion coefficient, respectively. The waist of the beam w_0_ was measured prior to each experiment using 100 nM solutions of eGFP and mCH in water assuming their diffusion coefficients are 90 µm²/s (65, 66). Finally, the diffusions maps were obtained by calculating for each pixel of the image, the average diffusion coefficient in a surrounding area of 64×64 pixels (10.24 µm²). In the resulting diffusion maps, the pixels are color coded by the average D value in the surrounding area.

## RESULTS

### Fluorescent labelling of HIV-1 gRNA and Gag proteins in cells

We transfected a stable HeLa cell line expressing MS2-fused to eGFP (here called HeLa MS2-eGFP) (23) with a plasmid encoding a modified HIV-1 gRNA (pIntro) containing a cassette of 12 MS2 stem-loops recognized by the MS2-eGFP protein (Figure 1A-1 and Figure 1A-3). Of note, in our system, the eGFP contains a nuclear localization signal (NLS) that directs MS2-eGFP towards the nuclei and nucleoli (Figure 1A-2). To fluorescently label Gag proteins, we fused the mCH probe upstream of the CA domain, in order to minimize the impact of the tag on protein activities (44, 63) (Figure 1B).

At first, we transfected the HeLa MS2-eGFP cells with the plasmids mentioned above and imaged them 24 h later. In cells transfected with pIntro, the phenotype observed was characterized with non-fluorescent nucleoli, in contrast to non-transfected cells where fluorescence was mainly concentrated at those sites (Figure 1A and 2A). Moreover, by using immunofluorescence with an antibody directed against the RNA polymerase II Phospho S2 (Figure S1), we observed that the bright green clusters in the nucleoplasm (Figure 2A, white arrows) corresponded to active transcription sites. In a further step, the co-transfection of pIntro with a Rev-encoding plasmid ensured the complete export of the MS2-eGFP-labeled gRNA from the nucleus to the cytoplasm due to the specific recognition of the RRE (Rev Response Element) sequence by Rev (Figure 2B). Finally, when unlabelled Gag was expressed alone (Figure 2C) or together with Gag-mCH (Figure 2D), the gRNA was re-localized to the PM. These observations indicated that the MS2-eGFP based strategy is well suited to investigate the interactions between HIV-1 gRNA and Gag proteins by fluorescence-based techniques.

**FIGURE 2:**
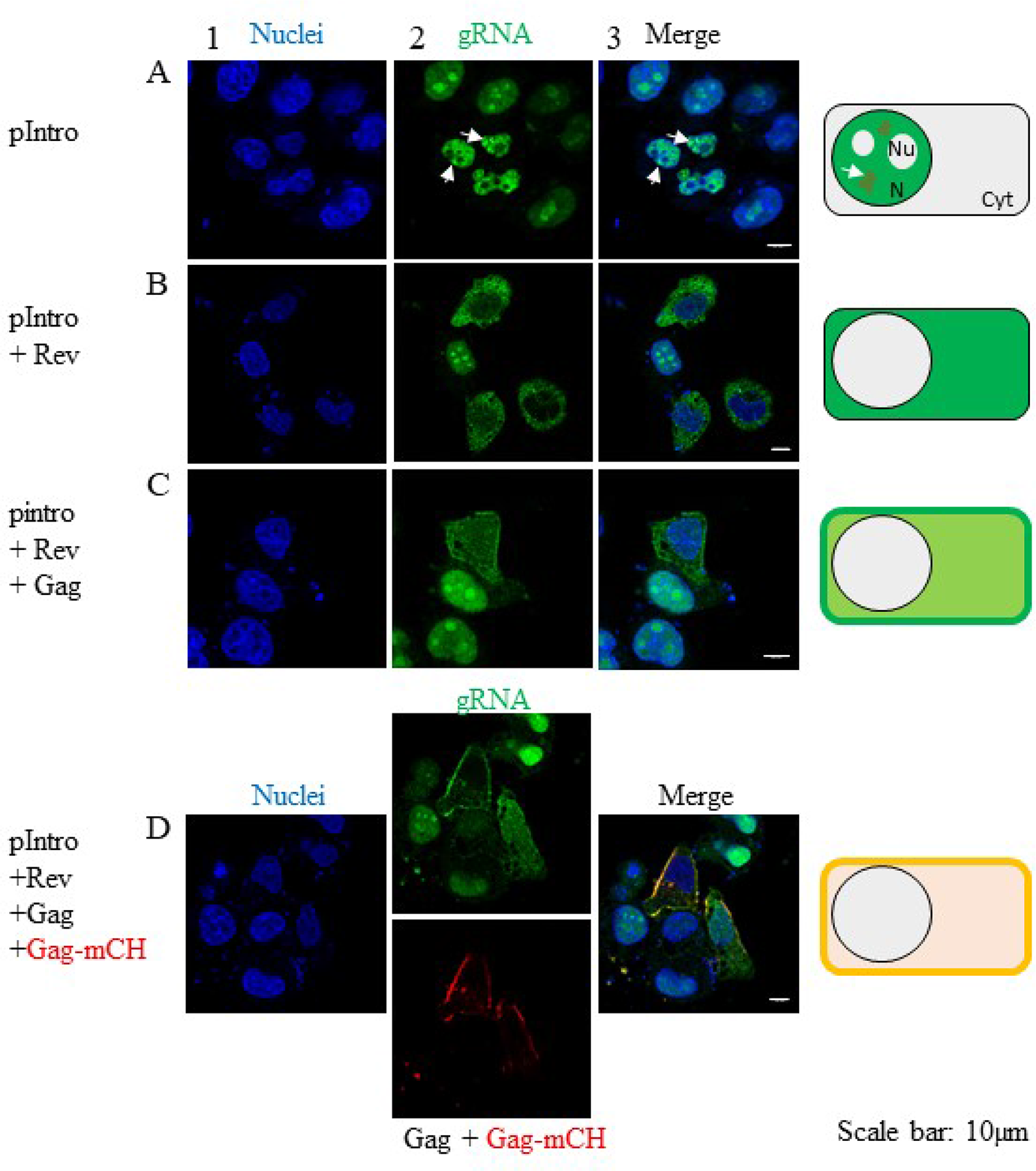
Membrane re-localization of HIV-1-gRNA by Gag. (A) HeLa cells stably expressing MS2-eGFP were transfected with a construct encoding p-Intro (see Materials and Methods). Nuclei were stained with Hoechst33258 (blue channel, column 1). Non-transfected cells showed GFP signal in the nuclei and nucleoli, while in cells transfected with pIntro, the MS2-eGFP fluorescence signal was only localized in the nuclei but no more in the nucleoli (green channel, column 2). The merge is in column 3. (B) The cells were then transfected with a plasmid encoding for Rev. This co-transfection ensured the complete export of the MS2-eGFP-labelled gRNA from the nucleus to the cytoplasm due to the specific recognition of the RRE. When Gag alone (C) or in mixture with Gag-mCH (D) was co-expressed, gRNA was re-localized to the PM. Confocal microscopy was performed 24 h post transfection. Cartoons on the right illustrate the observed localizations of MS2-eGFP-RNA. N (Nucleus), NU (Nucleolus), Cyt (Cytoplasm).

### At least one ZF of Gag is required for gRNA enrichment at the PM

To investigate the impact of mutations in the NC domain of Gag on the cellular localization of gRNA, we used a Gag mutant where the complete NC domain was deleted (GagΔNC), as well as Gag mutants carrying either a single (GagΔZF1 or GagΔZF2) or a double ZF deletion (ΔZF1-2) (Figure 1B). We also included a non-myristoyled Gag protein in which the Gly at position 2 was substituted with an Ala residue (GagG2A), thus preventing the addition of a myristate group (Figure 1B). Globally, we observed 24 h post transfection by confocal microscopy that all tested Gag proteins displayed a PM localization (Figure 3A, column 2), with the exception of the GagG2A mutant which was found exclusively in the cytoplasm, as expected (67) (Figure3A, column 2). A careful quantification (see Materials and Methods) showed that WT Gag and gRNA co-localized at the PM in 84 ± 3 % of cells, while this percentage dropped to 71 ± 3 % and 57 ± 1 % for GagΔZF1 and GagΔZF2, respectively (Figure 3A and 3B). Interestingly, gRNA was found to accumulate in the cytoplasm in presence of GagΔZF1-2 or GagΔNC, and no co-localization of the proteins with gRNA was observed at the PM in these cases (Figure 3A and 3B). Altogether, these experiments show that the two ZFs of the NC domain of Gag are required for an optimal trafficking of gRNA to the PM. However, the presence of one ZF is sufficient to partially relocate gRNA from the cytoplasm to the PM.

**FIGURE 3:**
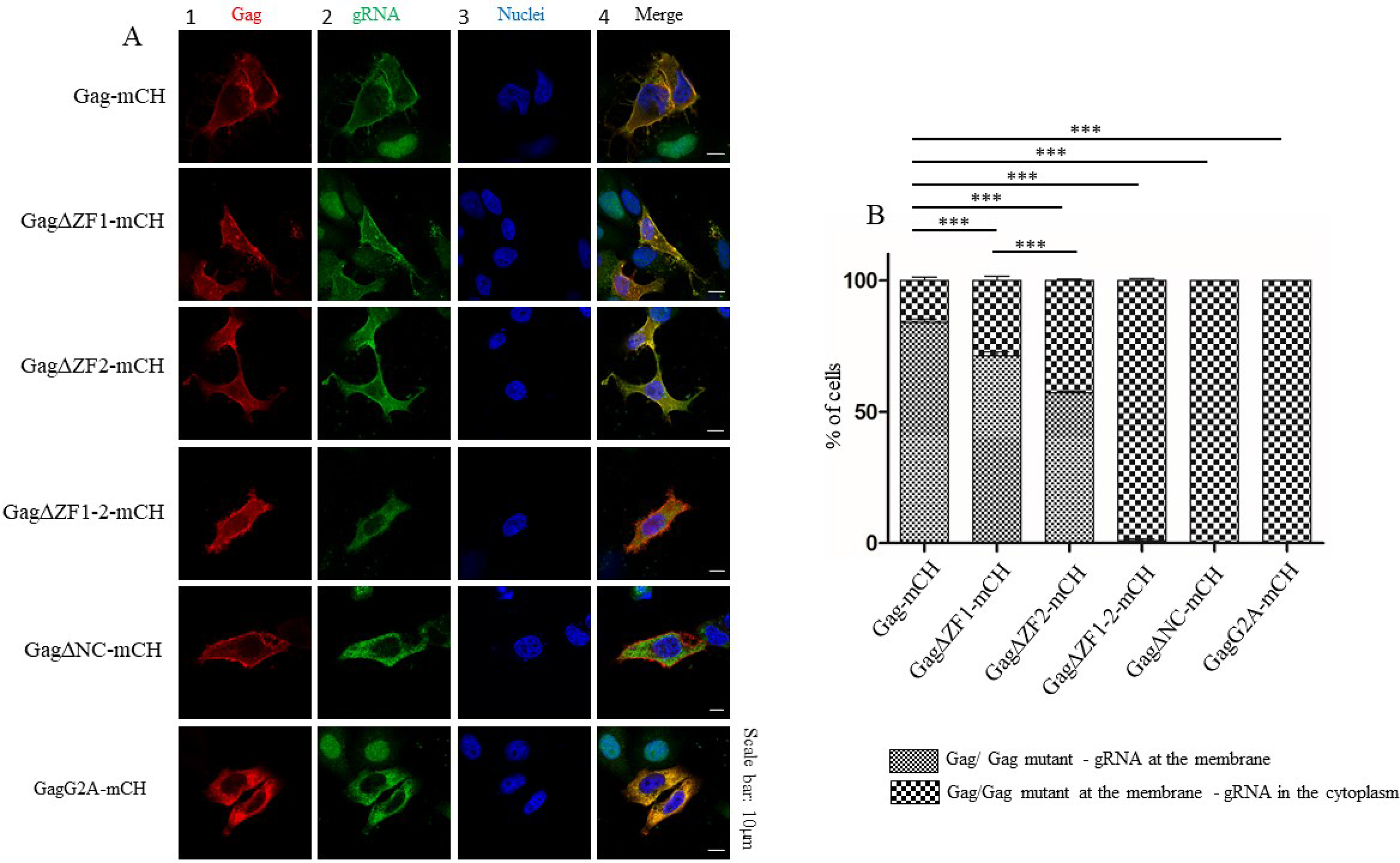
Confocal microscopy of MS2-eGFP HeLa cells co-expressing Gag-mCH proteins and gRNA. (A) The localization of Gag-mCH proteins (column 1, red channel) and MS2-eGFP-gRNA (column 2, green channel), as well as the staining with Hoechst33258 as a fluorescent marker for the nucleus (column 3, blue channel) and the merge of these images (column 4) are shown. Each panel indicates the major observed phenotype. (B) Histograms show the percentage of cells in which gRNA was found to diffuse in the cytoplasm (large dots), or alternatively was localized at the PM (small dots), in the presence of the different Gag-mCH proteins. Cells were imaged 24 h post-transfection by confocal microscopy. We counted 100 cells per condition. The analysis was performed on 4 independent experiments and error bars represent the standard error of the mean (SEM). Statistics was obtained with a χ^2^ test and revealed a significant difference (*** p <0.001). The scale bar of 10 μm is indicated.

### Real-time kinetics of gRNA co-accumulation with Gag at the PM

Next, we performed two-color time-lapse microscopy experiments to monitor in real time the events taking place between gRNA transcription and its localization at the PM in living cells. To this aim, HeLa MS2-eGFP expressing cells were microinjected with a combination of plasmids expressing gRNA, Gag and Rev and imaged every 5 min for 4 h. About 5 min after microinjection, the MS2-eGFP fluorescence accumulated as clusters in the nucleoplasm corresponding to active transcription sites (Figure S1), while the nucleoli appeared non fluorescent (Movie S1). When the viral Rev factor was expressed, the MS2-eGFP labelled gRNA was then found to accumulate at the nuclear envelope, and to colocalize with the nuclear envelope marker POM121-mCH (Figure S1B and Movie S1). The Rev-driven export of gRNA was subsequently observed through the green fluorescence signal accumulating in the cytoplasm. About 1 h after microinjection, Gag-mCH appeared in the cytoplasm, and we evaluated the average delay between the appearance of the Gag-mCH proteins in the cytoplasm and the accumulation of the first MS2-eGFP labeled gRNAs at the PM (Figure 4A and 4B). In agreement with the conclusions of the previous paragraph, we observed that the enrichment of the MS2-eGFP labelled gRNA at the PM after 4 h was observable for less than 7% of cells expressing GagΔZF1-2 (Movie S2) or GagΔNC (Movie S3). For cells expressing GagΔZF1(Movie S4) and GagΔZF2 (Movie S5), we noticed gRNA accumulation to the PM, but with a significant delay as compared to WT Gag proteins. Indeed, while gRNA accumulated at the PM within 47 ± 4 min (n = 39) in presence of WT Gag, it took about 73.5 ± 4 min (n=39) and 94.5 ± 5 min (n=50) in the case of GagΔZF1 and GagΔZF2, respectively (Figure 4B). In a further step, we monitored the mean delay between Gag-mCH appearance at the PM and the accumulation of MS2-eGFP labeled gRNAs at the same sites. Similarly, to our previous observation, GagΔZF2 showed a significantly increased delay 45 ± 3 min (n=50) compared to WT Gag 17 ± 3 min (n=26) or to GagΔZF1 23.5 ± 5 min (n=39) (Figure 4C). These results suggest that deletion of the ZF motifs in Gag introduces a delay in the co-localization of gRNA at the PM, and that ZF2 has a greater impact than ZF1 in the incorporation of gRNA into Gag clusters at the PM assembly sites.

**FIGURE 4:**
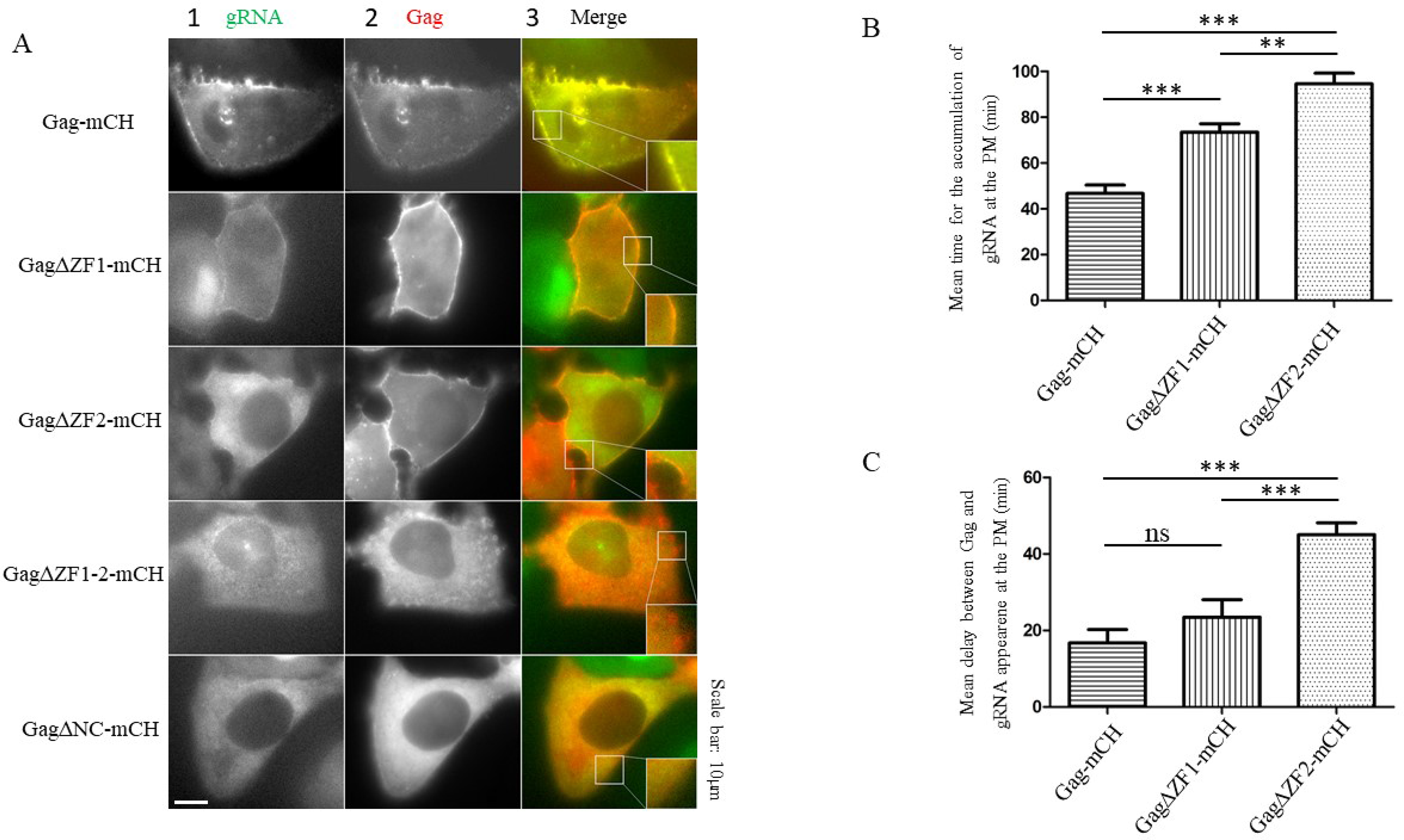
**Kinetic analysis of gRNA localization at the PM induced by Gag proteins**. (A) Time-lapse microscopy on cells microinjected with a combination of plasmids expressing MS2-eGFP-gRNA (column 1, green channel), Gag-mCH (column 2, red channel) and Rev and imaged every 5 minutes for 4 h. In the merge colum 3, the insets correspond to a zoom of the PM region. The scale bar of 10 μm is indicated. (B) Quantification of the average delay separating the appearance of Gag-mCH proteins in the cytoplasm and the detection of the first MS2-eGFP-gRNA accumulating at the PM. (C) Quantification of the average delay separating the detection of Gag-mCH and MS2-eGFP-gRNA signals at the PM. The statistical analysis was performed by one-way ANOVA associated to Tukey’s multiple comparison tests and revealed significant differences (2 stars p<0.01, 3 stars p<0.001, error bars represent Standard Errors) between Gag, GagΔZF1 and GagΔZF2 (26-50 cells analyzed per experiment).

### Monitoring the interactions between Gag proteins and gRNA at the PM

To further demonstrate the direct interaction between Gag and gRNA at the PM, we performed FRET-FLIM (Fluorescence Lifetime Imaging Microscopy). FRET occurs when the FRET donor (eGFP linked to MS2) and acceptor (mCH bound to Gag) are less than 10 nm apart. The FLIM technique is based on the analysis of the donor lifetime at each pixel of the image. When FRET occurs, the donor lifetime decreases. Of note, the lifetime is independent of the local concentration of fluorophores and the instrumental setup. Typically, FLIM images are built up using a false color scale covering the range of donor lifetimes from 2 ns (red) to 2.4 ns (blue). This allows a direct description of each pixel in terms of FRET efficiency and thus About 24 h after transfection of MS2-eGFP HeLa cells, the lifetime value of MS2-eGFP-gRNA in the presence of the mCH acceptor alone was similar to the lifetime of MS2-eGFP in the nuclei of non-transfected cells (about 2.3 ns). This reflects the absence of FRET between the probes under these conditions, and the fact that the fluorescence lifetime of MS2-eGFP is not influenced by its binding to gRNA. In the presence of Gag-mCH proteins, we observed a decrease of the lifetime of the MS2-eGFP-gRNA complexes at the PM (Figure 5A), demonstrating that FRET occurs between Gag and gRNA at those sites. According to Eq.1 (see Materials and Methods), the corresponding value for FRET efficiency was 5 ± 0.5 %. (Figure 5B). In cells transfected with the mCH-labeled GagΔZF1 or GagΔZF2, FRET efficiency was about 8 ± 1% and 9 ± 2 %, respectively (Figure 5A panels 3-4, and Figure 5B). FLIM-FRET analysis thus confirmed that Gag proteins and gRNA interact at PM and the deletion of one ZF does not affect the interaction of Gag with gRNA at these sites.

**FIGURE 5:**
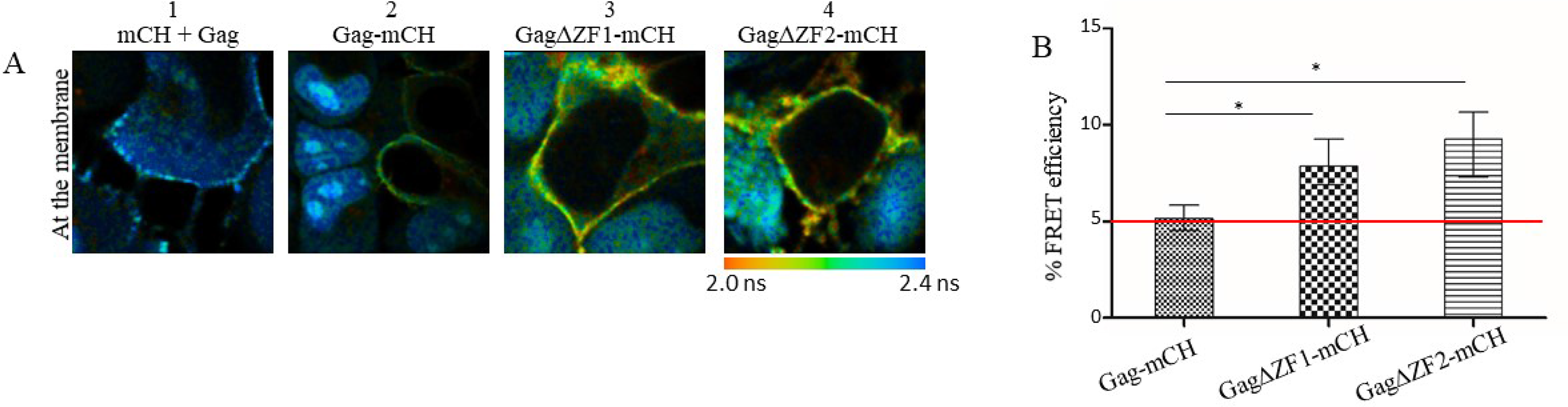
**FRET-FLIM analysis of the interaction between gRNA and Gag at the PM**. (A) MS2-eGFP HeLa cells were transfected with a combination of plasmids and FLIM analysis in the cytoplasm was carried out 24 h post transfection. The fluorescence lifetime of MS2-eGFP-gRNA was determined by using a single exponential model and was color coded, ranging from red (2.0 ns) to blue (2.4 ns). FLIM images of gRNA in the presence of unlabeled Gag and free mCH [1], Gag-mCH [2], GagΔZF1-mCH [3], or GagΔZF2-mCH [4]. (B) Corresponding histograms representing FRET efficiencies for Gag-mCH (small dots), GagΔZF1-mCH (large dots) and GagΔZF2-mCH (lines). We performed three independent experiments on at least 30 cells. The red line indicates the threshold value (5%) above which FRET efficiency can be considered as corresponding to a direct interaction between fluorescently labelled gRNA and Gag proteins (44). FRET efficiency values were calculated as described in Materials and Methods (Eq.1). The statistical analysis was realized by a Student’s T-test with significant differences represented by 1 star * p<0.05. All images were acquired using a 50 μm×50 μm scale and 128 pixels × 128 pixels.

### Monitoring the interaction between Gag proteins and gRNA in the cytoplasm

We then investigated the interaction between Gag and gRNA in the cytoplasm. We imaged by FRET-FLIM the cells 16 h after transfection, when large quantities of Gag proteins are still present in the cytoplasm. Interestingly, the expression of Gag-, GagΔZF1- and GagΔZF2-mCH proteins led to a decrease of MS2-eGFP/gRNA (the donor) lifetime in the cytoplasm, as can be seen from the colour change from blue (Figure 6A, panel 1) to green (Figure 6A, panels 2-4). The corresponding FRET efficiency values were of 6.6 ± 0.8 %, 6.3 ± 0.2 %, and 6.4 ± 1 %, respectively, indicating that these proteins interact with the gRNA in the cytosol. In contrast, FRET efficiencies for GagΔZF1-2-mCH and GagΔNC-mCH were only 1.3 ± 0.9 % and 1.6 ±1 %, respectively, suggesting that one ZF motif is necessary and sufficient for the interaction between Gag and gRNA in the cytoplasm (Figure 6A, panels 5-6). Finally, the non-myristoylated Gag mutant (GagG2A-mCH) was also found to interact with gRNA in the cytoplasm, with a FRET efficiency of 7.8 ± 0.1 % (Figure 6A, panel 7) indicating that myristoylation is not necessary for Gag-gRNA interaction in the cytosol.

**FIGURE 6:**
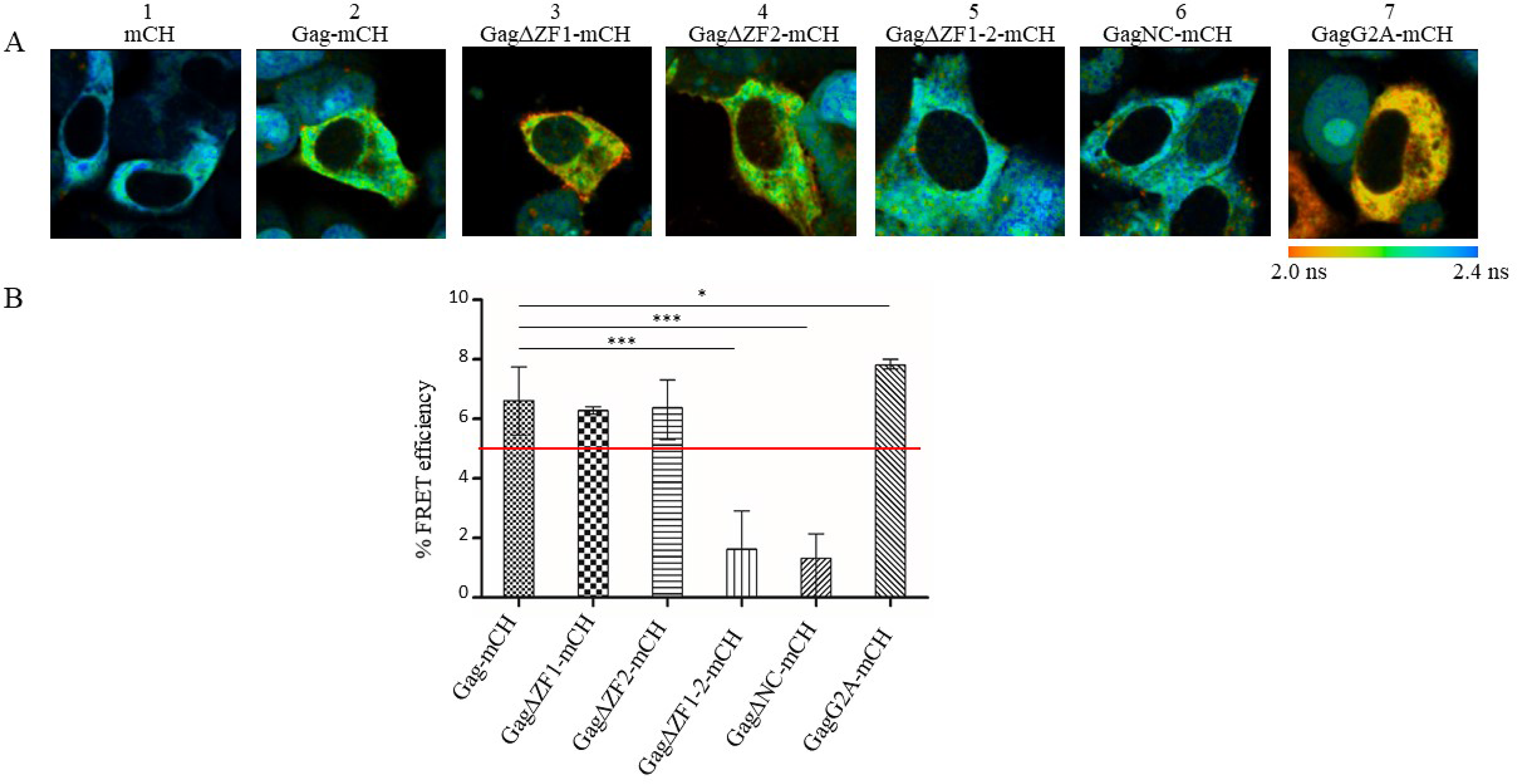
**FRET-FLIM analysis of the interaction between gRNA and Gag in the cytoplasm**. (A) MS2-eGFP HeLa cells were transfected with our combination of plasmids and FLIM analysis in the cytoplasm was carried out 16 h post transfection. The fluorescence lifetime of MS2-eGFP was determined by using a single exponential model and was color coded, ranging from red (2.0 ns) to blue (2.4 ns). FLIM images of gRNA in the presence of unlabeled Gag and free mCH [1], Gag-mCH [2], GagΔZF1-mCH [3], GagΔZF2-mCH [4], GagΔZF1-2-mCH [5], GagΔNC-mCH [6], or GagG2A-mCH [7]. (B) Corresponding histograms represent FRET efficiencies for Gag and mCH (small dots), Gag-mCH (large dots), GagΔZF1-mCH (horizontal lines), GagΔZF2-mCH (vertical lines), GagΔZF1-2-mCH (vertical lines), GagΔNC-mCH (oblique lines up), or GagG2A-mCH (oblique lines down). The bars represent the means ± standard error of the mean for three independent experiments on at least 30 cells. The red line indicates the threshold value (5%) above which FRET efficiency values can be considered as corresponding to a direct interaction between fluorescently labelled gRNA and Gag proteins. The statistical analysis was realized by a Student’s T-test with significant differences represented by 1 star * p<0.05, 2* p<0.01 and 3* p<0.001. All images were acquired using a 50 μm×50 μm scale and 128 pixels × 128 pixels.

Next, the cytoplasmic diffusion of Gag and gRNA was investigated by Raster Image Correlation Spectroscopy (RICS) (65, 68, 69). This method is based on the analysis of the fluorescence intensity fluctuations between neighboring pixels by spatially autocorrelating the image in x and y directions. The resulting spatial correlation surface (SCS) is fitted with a 3D diffusion model to obtain the value of the diffusion coefficient (D) of the macromolecules in the scanned area. In a first experiment, we measured the cytoplasmic diffusion of MS2-eGFP. Stacks of 50 images were recorded in the cytoplasm of living cells (Figure 7A red frame, and 7B), and the mean SCS were calculated (Figure 7C). As a result of the presence of the NLS sequence, the majority of the MS2-eGFP molecules were located in the nucleus (Figure 7A) even though, a fraction of the MS2-eGFP molecules (∼25-30% based on RICS and intensity fluorescence measurements) was found to diffuse in the cytoplasm. The average diffusion coefficient of the cytoplasmic MS2-eGFP molecules was 1.8 µm/s² (Figure 7D, white bar), which is not consistent with the theoretical estimation based on the size of MS2-eGFP construct. Indeed, the hydrodynamic radius of the MS2-eGFP protein r_MS2-eGFP,_ (calculated as described in (70)) is 3.35 nm, and this value is 1.12 fold larger than the hydrodynamic radius (R_h_) of eGFP alone (reGFP =2.8 nm, (71)). Since the diffusion coefficient is inversely proportional to the radius of the diffusing molecule, a D value of 16.7 µm²/s for the free MS2-eGFP is expected from the ratio of the r_eGFP_/r_MS2-eGFP_ and the previously determined D_eGFP_ value (∼20 µm²/s, (72)). The comparison between the expected and experimental D values strongly suggests that MS2 protein may bind to cellular factors, likely cellular RNAs in the cytoplasm. Besides, in the absence of pIntro, expression of Gag-mCH or GagΔNC-mCH did not affect the diffusion of MS2-eGFP (Figure 7D). In contrast, the expression of pIntro and Rev led to a strong decrease in the D value, likely as the result of the binding of MS2-eGFP to viral gRNA and its subsequent relocation from the nucleus to the cytoplasm (Figure 7A, yellow frame). The mean value of D for MS2-eGFP bound to gRNA was about 0.3 µm²/s, in line with previous analysis on HIV-1 RNA diffusion by tracking assays (73). Furthermore, the interaction with Gag-mCH, or GagΔNC-mCH proteins did not significantly affect the diffusion of HIV-1 gRNA, which is consistent with the binding of a limited number of Gag copies to gRNA (27).

**FIGURE 7:**
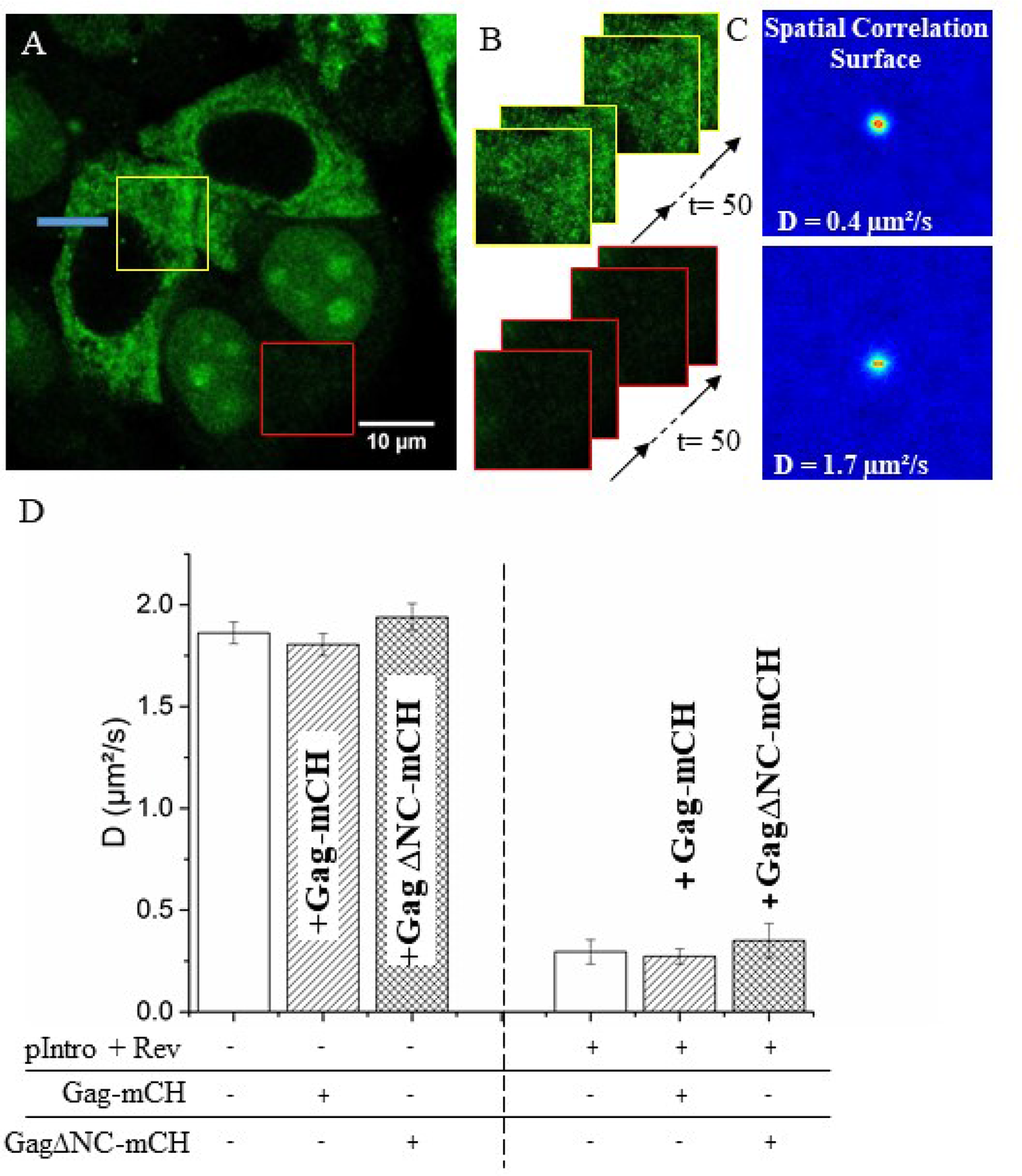
**RICS analysis of MS2-eGFP diffusion in the cytoplasm**: (A) Confocal images of MS2-eGFP expressing cells transfected with plasmids expressing pIntro and Rev. For the RICS measurements, stacks of 50 images were recorded in the cytoplasm (B) and the average Spatial Correlation Surfaces were then calculated (C) and fitted with a 3D free diffusion model. (D) Diffusion coefficient values of MS2-eGFP in cells expressing or not expressing pIntro and Rev (white), Gag-mCH (oblique lines) and GagΔNC-mCH (dashed). The presence of pIntro and Rev (right panel) induces a drastic decrease of the MS2-eGFP diffusion coefficient (left panel). For each condition, the measured values represent the mean and the corresponding (SEM) of 50-60 measurements in three independent experiments (15 - 20 cells analyzed per experiment). The statistical analysis was realized by a Student’s T-test with significant differences represented by 1 star * p<0.05 2* p<0.01 and 3* p<0.001.

In a next step, we monitored by RICS the cytoplasmic diffusion of Gag-mCH. Since the size of Gag is significantly smaller than the size of gRNA, the association of Gag proteins with gRNA should produce a large decrease in the value of their diffusion coefficient. The measurements were performed in a cytoplasmic volume in the midplane of the cell (Figure 8A). The focal planes of the acquisition were chosen carefully to minimize possible artifacts due to Gag-mCH molecules bound to the PM. The mean D value for Gag in the absence of gRNA was 1.1 ± 0.6 µm²/s (Figure 9B), in reasonable agreement with previous reported D values of 2.4 ± 0.5 µm²/s (74, 75). These values are considerably smaller than the theoretical value (13.8 µm^2^/s) calculated assuming that the molecular weight of Gag-mCH is about 82 kDa and using an empirical formula relating the molecular weight to the R_h_ (70). The discrepancy between the theoretical and experimental values is likely due to Gag-mCH tendency to multimerize and interact in the cytosol with cellular factors. This conclusion was further strengthened by the D value of 10.4 ± 2.1 µm^2^/s measured for the cytoplasmic diffusion of eGFP trimers (Mw = 81 kDa) (in good agreement with 9.5 µm²/s reported by (74)) that have approximately the same size as Gag-mCH and are not supposed to bind to any cellular components (44, 76, 77). In line with a previous study (77), mutations affecting the myristoylation site do not affect Gag mobility (D_GagG2A_ = 1.3 ± 0.6 µm²/s). Interestingly, D values of 1.7 ± 0.6 µm²/s and 1.6 ± 0.5 µm²/s were obtained for GagΔZF1 and GagΔZF2, respectively, while deletion of the two ZFs or the complete NC domain resulted in increased D values of 2.3 ± 0.9 µm²/s and 4.7 ± 1 µm²/s. These high D values (77) are likely related to the inability of GagΔZF1-2 and GagΔNC to multimerize and bind to cellular RNAs and proteins (44, 76). We then performed the same analysis in cells expressing HIV-1 gRNA. The diffusion coefficients of Gag-mCH, GagG2A, GagΔZF1 and GagΔZF2 proteins decreased significantly (∼25-30%) in the presence of gRNA (Figure 8B, grey bars), while no effect was observed for GagΔNC and GagΔZF1-2 mutants.

**FIGURE 8:**
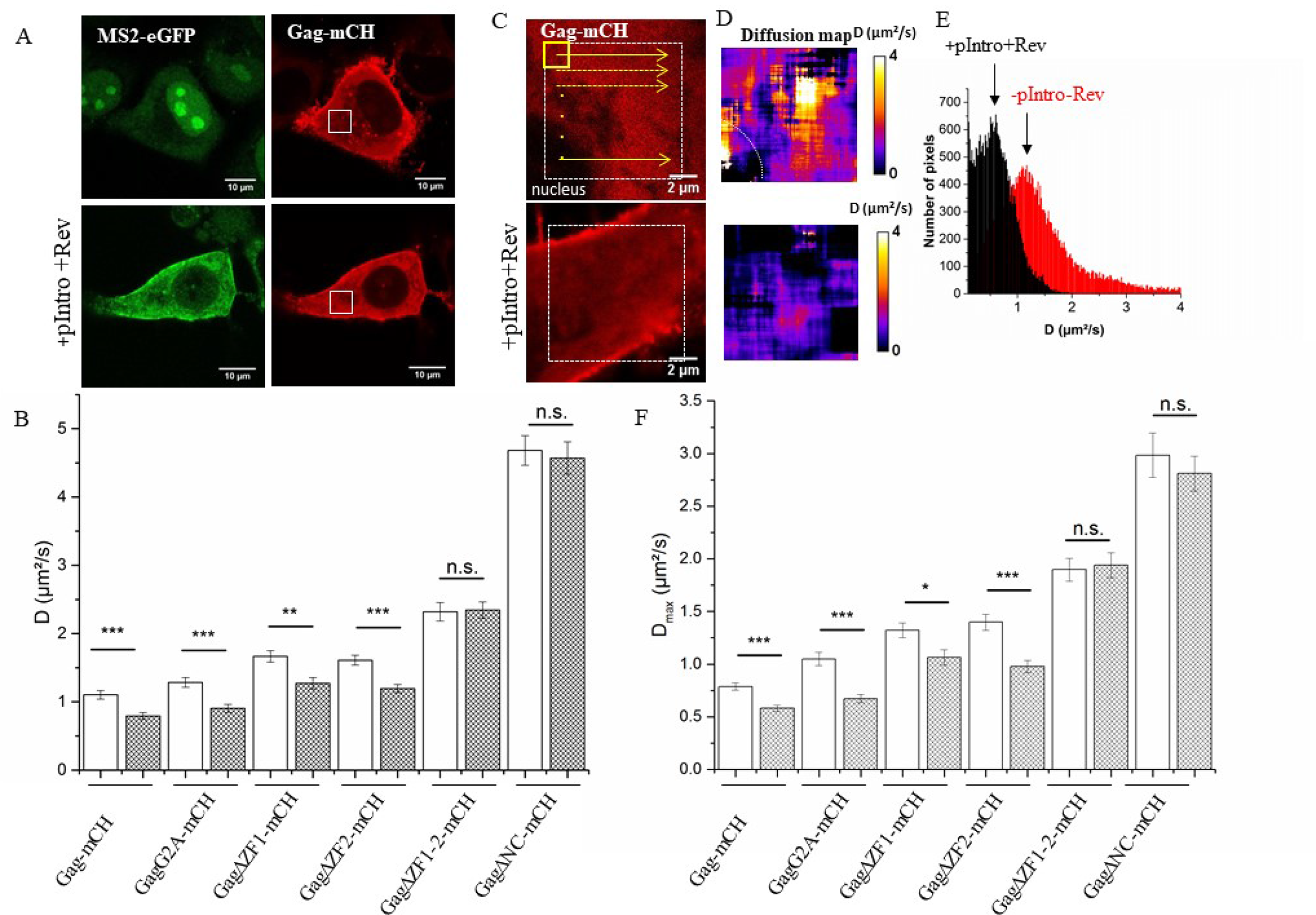
**RICS analysis of Gag-mCH diffusion in the cytoplasm**. (A) Confocal images of MS2-eGFP and Gag-mCH expressing cells in the absence (top panels) and the presence (bottom panels) of pIntro and Rev. MS2-eGFP-gRNA was observed in the cell cytoplasm (green channel) and the RICS measurements were performed on the labeled Gag proteins in the red channel. (B) Diffusion coefficient values of Gag proteins in the presence (dashed bars) and the absence (white bars) of pIntro and Rev. (C) Confocal images and (D) corresponding diffusion maps of Gag-mCH in the absence (top panels) and the presence (bottom panels) of pIntro and Rev. (E) Histogram representation of the D values of the diffusion maps. The arrows show the positions of the most frequent D values, called D_max_. (F) D_max_ values of Gag proteins in the presence (dashed) and absence (white) of pIntro and Rev. In (B) and (D), the measured values represent the mean and the corresponding SEM of 50-60 measurements in three independent experiments (15-20 cells analyzed per experiment). The statistical analysis was realized by a Student’s T-test with significant differences represented by 1 star * p<0.05 2* p<0.01 and 3* p<0.001.

To further strengthen our analysis, we mapped the diffusion coefficients of the Gag proteins in a larger part of the cell. To this aim, a window of 64×64 pixels was shifted pixel by pixel along the images and an average D value was calculated for each position (Figure 8C). A diffusion map was then generated by representing the D values obtained in each pixel. Examples of diffusion maps of Gag-mCH proteins in the presence and absence of gRNA are shown in Figure 8D. The histogram of the diffusion maps (Figure 8E) revealed that the D values are highly variable, and significantly decreased in the presence of gRNA. Comparison of the D values at the maximum of the histograms (D_max_) indicated that in good agreement with our analysis (Figure 8B), the D_max_ value decreased by 20-35% for Gag-, GagG2A-, GagΔZF1- and GagΔZF2-mCH proteins, while the D_max_ value remained constant for GagΔZF1-2- and GagΔNC-mCH mutants (Figure 8F). Thus, the RICS data confirmed that Gag, GagG2A, and Gag mutants carrying only one zinc finger deletion bind to the gRNA, while GagΔZF1-2- and GagΔNC mutants do not, in agreement with our FRET/FLIM conclusions (Figure 6).

## DISCUSSION

Previous biochemical and genetic studies have extensively investigated *in vitro* the role of the two ZFs of NCp7 (56, 57, 59 –62, 78–83). In this study, we combined several imaging techniques to obtain a clear picture of the role of both ZFs in the NC domain of Gag in the intracellular trafficking of HIV-1 gRNA to the PM assembly sites. Even though cellular Gag-gRNA interactions were already described (18, 74), our analysis focused on the comparison of the gRNA interactions with WT Gag and Gag mutants where either one or both ZFs were deleted. Our analysis evidenced that at least one ZF is required for an efficient interaction between Gag and gRNA in the cytoplasm and at the PM. While the two ZFs seem to be redundant for this interaction, ZF2 played a more important role than ZF1 in the trafficking of the ribonucleoprotein complexes to the PM (Figure 5, 6 and 8).

By performing real-time analysis in the presence of Gag, we found that gRNA accumulated at the PM within 46.7 ± 3.7 min in presence of Gag (Figure 4A and 4B). This result is in good agreement with previous studies showing an accumulation of gRNA dimers at the PM during virus assembly in about 30 min (3, 21, 84). On the other hand, the deletion of a single ZF delayed the gRNA accumulation at the assembly sites, as it took about 73.5 ± 3.7 min and 94.5 ± 4.7 min to accumulate gRNA at the PM in the case of GagΔZF1 and GagΔZF2, respectively (Figure 4B). Data from the literature showed that the NC domain and the C-terminal p6 domain in Gag are both involved in the budding cellular machinery, since deletions of the NC domain or its two ZFs were found to interfere with virus release by impairing the recruitment of Tsg101 ESCRT-I proteins and their co-factors, such as ALIX (44, 85, 86). Moreover, it was reported that the deletion of the distal ZF2 led not only to abnormal uptake of Tsg101, but also to biogenesis defects during virion formation (83). It is thus possible that the observed delay of GagΔZF1 or GagΔZF2 to reach the PM (Figure 4) is related to their defective recruitment of the ESCRT machinery.

In a further step, the non-equivalence of the two ZFs in viral RNA recruitment to the PM was confirmed by the mean delay observed between Gag-mCH appearance at the PM, and the accumulation of MS2-eGFP labeled gRNAs at the same sites (Figure 4C). Indeed, GagΔZF2 showed a significantly increased delay 45 ± 3 min compared to WT Gag 17 ± 3 min or to GagΔZF1 23.5 ± 5 min (Figure 4C), confirming that ZF2 has a greater impact than ZF1 in the recruitment of gRNA to the PM. Thus, even though the two ZFs displayed redundant roles in the cytoplasmic context, we observed that ZF2 played a more prominent role in the trafficking of the gRNA/Gag complexes to the assembly sites at the PM. The idea that the two ZF these ZFs do not seem to be functionally equivalents was also supported by recent *in vitro* data in the NCp7 context showing that the ZF2 would regulate of association with NAs, because of its larger accessibility compared to ZF1 which would contribute to stabilize the resulting complex (62).

Our RICS analysis further showed that Gag proteins did not affect the diffusion of HIV-1 gRNA (Figure 8), likely because of the limited size-increase of the gRNA upon binding of a few Gag proteins. This is fully consistent with the notion that Gag multimerization could be initiated in the cytoplasm, and then triggered by RNA binding, (26, 75, 76, 87–89), and with our previous *in vitro* data showing that a limited number of Gag proteins (i.e. about two trimers) bind to gRNA fragments (14). Deletion of the NC domain induced a significant increase in diffusion compared to WT Gag (4.7 ± 1 µm²/s *vs* 1.1 ± 0.6 µm²/s), in line with previous data on Gag mutants in which all basic residues of the NC domain were replaced by Ala residues (77). This increased diffusion could be explained by the impacted capacity of Gag to multimerize and to bind to cellular factors when its NC domain is deleted (44, 76).

Our data also included the G2A mutant, where the absence of myristate does not only abolish the anchorage of Gag at the PM, but also impacts Gag oligomerization (76, 90). In good agreement with the literature (77), our findings showed that mutations affecting myristoylation did not affect Gag mobility, nor cytosolic binding to gRNA. This definitely supports the conclusion that, the binding of HIV-1 Gag to viral RNA and to PM are independent events governed by different domains (Figure 3 and 6). However, how MA and NC domains are employed by retroviral Gag to interact with RNA is retrovirus-specific, since previous observations on deltaretrovirus showed that HTLV-2 MA has a more robust chaperon function than HTLV-2 NC and contributes importantly to the gRNA packaging (91).

Altogether our findings show for the first time that the two zinc fingers in the nucleocapsid domain of the HIV-1 Gag precursor are equivalent for the interaction with the genomic RNA in the cytoplasm, and ZF2 has a more important role than ZF1 for the intracellular trafficking of the ribonucleoprotein complex to the PM. Our data thus contribute to the current understanding and knowledge of the determinants governing the HIV-1 gRNA cellular trafficking to the assembly sites at the PM.

### Authors contributions

SB and HdeR designed the project. EB, SB and HdeR managed the project and drafted the manuscript with some assistance from the other co-authors. JCP, RM, and YM contributed to scientific discussions and to revise the manuscript. MbN, EB, PD and JB characterized the interactions between fluorescently labelled gRNA and Gag (confocal, FRET-FLIM, statistics). ER performed cloning. DD, RC and EB microinjected and imaged the dynamics of the interactions. HA performed RICS experiments; HA and PC performed the analysis of RICS experiments.

## Acknowledgments

The ANRS supported SB, JB and HdeR. We thank Romain Vauchelles for his assistance at the PIQ platform; Julien Godet and Frédéric Przybilla for their help in statistical analysis.

